## Supplementary information for "Structural basis for how sMAC is packaged for clearance"

### ONLINE SUPPLEMENTARY INFORMTATION

### SUPPLEMENTARY TABLES

#### Supplementary Table 1. CryoEM data collection and model validation statistics

##### Data collection specifications

| | Dataset-1 | Dataset-2 |
|---|---|---|
| **Microscope** | Titan Krios | Titan Krios |
| **Acceleration Voltage (keV)** | 300 | 300 |
| **Camera** | K2 | K2 |
| **Collection Mode** | Counting | Counting |
| **Pixel size (Å)** | 1.047 | 1.048 |
| **Stage tilt** | 0° | 37° |
| **# Frames** | 40 | 40 |
| **Integration time (s)** | 8 | 11 |
| **Total dose (e-/Å$^2$)** | 40 | 41.7 |
| **Defocus range (μm)** | -1.1 to -2.3 | -1.1 to -2.1 |
| **# Micrographs** | 11107 | 2596 |

##### Model validation statistics

| Structure | 2C9-sMAC | 3C9-sMAC |
|---|---|---|
| **Accession code** | PDB: 7NYD<br>EMD-12651 | PDB: 7NYC<br>EMD-12650 |
| **Map resolution (Å) - FSC threshold 0.143** | 3.3 | 3.5 |
| **# Residues** | 4940 | 5312 |
| **# Non-hydrogen atoms** | 38488 | 41373 |
| **Ramachandran outliers (%)** | 0.02 | 0.02 |
| **C-B outliers (%)** | 0.09 | 0.75 |
| **Clash score** | 2.20 | 3.63 |
| **Molprobity score** | 1.33 | 1.49 |
| **Map-Model FSC - threshold 0.5** | 3.2 | 3.5 |

#### SUPPLEMENTARY FIGURES

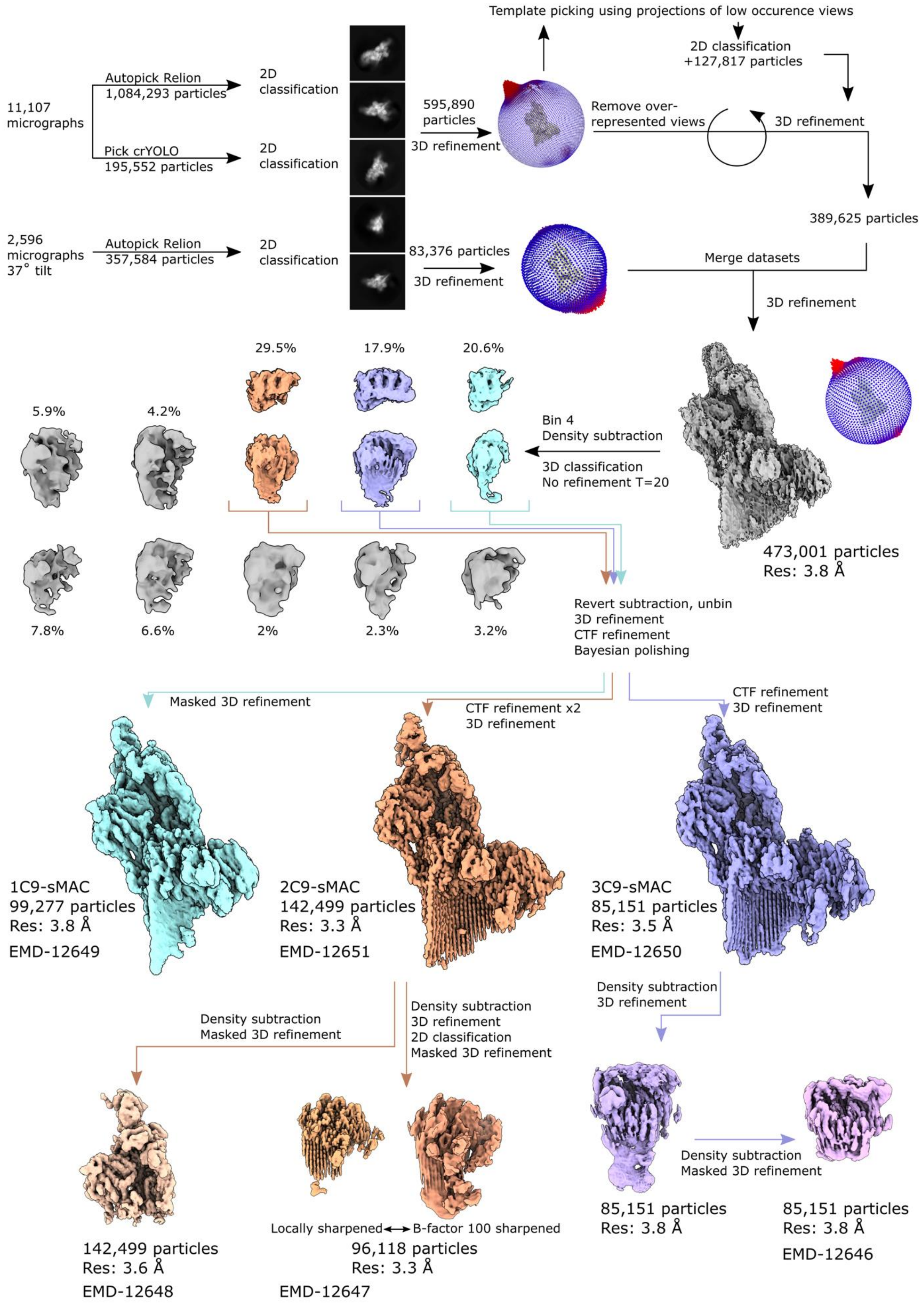

**Supplementary Figure 1:** CryoEM image processing workflow. Particles were picked from micrographs using autopicking programs in Relion and cryYOLO to account for differences in ice-thickness across the micrographs. Images were subject to 2D classification and an initial auto-refinement, resulting in a reconstruction with strong preferred orientations. To improve the angular distribution of the dataset, we combined three strategies: 1) We implemented template-based picking procedures using projections of low occurrence views. 2) We pruned particles from over-represented views. 3) We incorporated an additional dataset collected at a tilt angle of 37°. Duplicate particles were removed and datasets were merged. Data was subject to an additional 3D auto-refinement to generate a consensus sMAC reconstruction with an improved angular distribution. To separate sMAC maps that contained different numbers of C9 molecules we subtracted density corresponding to the core complement complex (C5b6, C7, C8) and used 3D classification with no refinement. Maps with clear density for either 1 (cyan), 2 (orange) or 3 (purple) copies of C9 were taken forward. Reconstructions for each of these three classes were then calculated based on the corresponding particles before density subtraction. Data was subjected to Bayesian polishing and multiple rounds of per-particle CTF refinement before a final 3D auto-refinement to generate the sMAC maps: 1C9-sMAC (3.8 Å) (EMD-12649), 2C9-sMAC (3.3 Å) (EMD-12651), and 3C9-sMAC (3.5 Å) (EMD-12650). To better resolve density above the C9 LDL domains, we used particles corresponding to the 3C9-sMAC map (purple) and subtracted density for the core complement complex. The resulting reconstruction (light purple) showed improved density in this region. We next performed an additional density subtraction and focused our 3D refinement on the core of C9 to generate the C9-clusterin focus-refined 3C9-sMAC map (pink, 3.8 Å) (EMD-12646). To better resolve density corresponding to C5b, we used particles corresponding to the 2C9-sMAC map (orange) and subtracted density for the MACPF arc. Using a masked 3D auto-refinement we calculated a map corresponding to C5b and C6/C7 C-terminal domains (tan, 3.6 Å) (EMD-12648). To better resolve the alternative conformation of C9 in sMAC, we used particles corresponding to the 2C9-sMAC map (orange) and subtracted density for the core complement complex. Particles were subjected to a further 2D classification to improve homogeneity of the population and a final reconstruction of the C9 focus-refined map was calculated (light orange, 3.3 Å) (EMD-12647). The locally sharpened map was used to build the intermediate C9 conformation in sMAC. Blurring the map using an ad hoc B-factor (100), we observe density in the vicinity of the hydrophobic β-hairpins of the C9 MACPF, which may correspond to vitronectin.

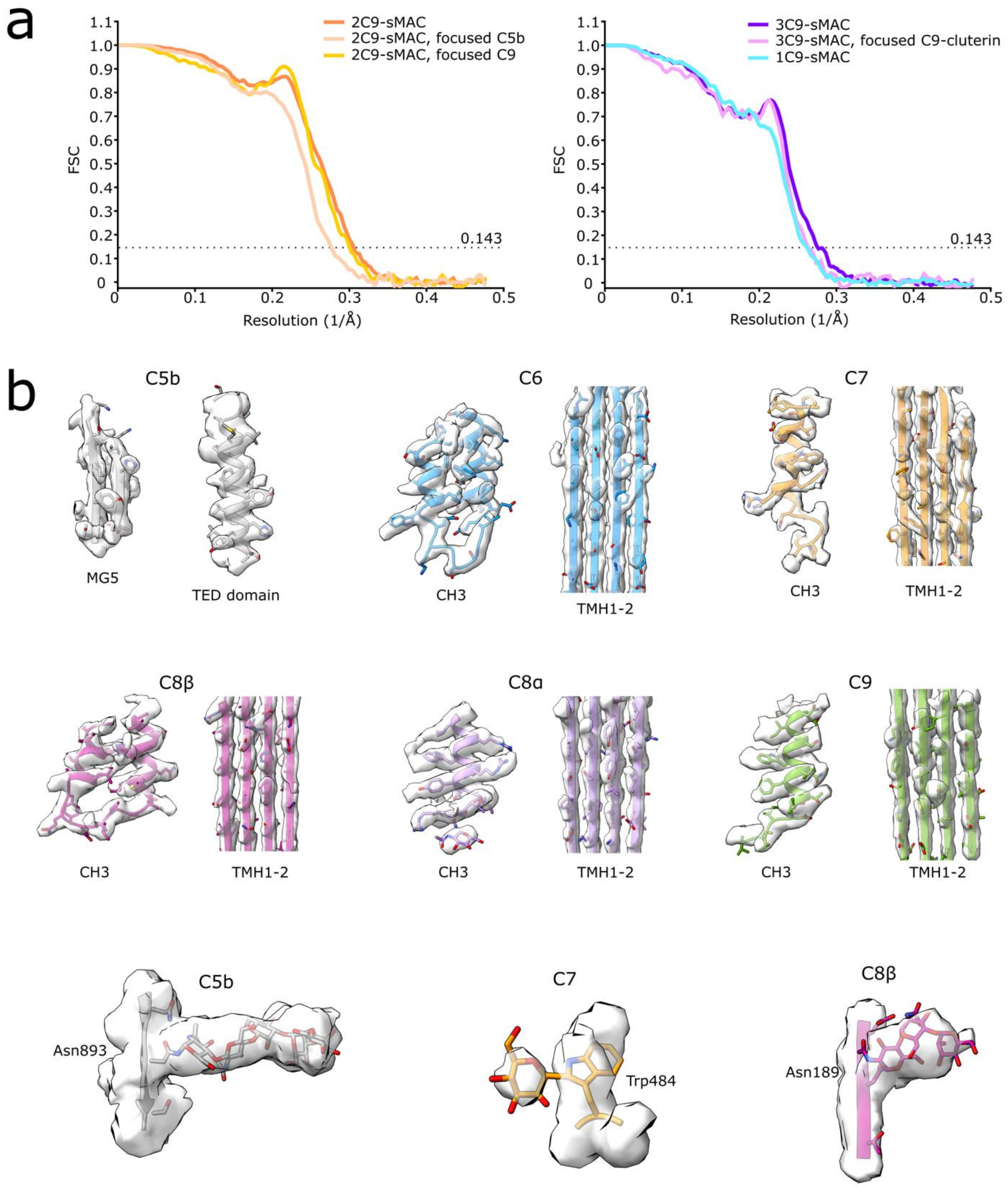


**Supplementary Figure 2:** CryoEM map validation. (a) Fourier Shell Correlation (FSC) curves for all deposited maps: 1C9-sMAC (EMD-12649), 2C9-sMAC (EMD-12651), 3C9-sMAC (EMD-12650), C5b-focused 2C9-sMAC (EMD-12648), C9-focused 2C9-sMAC (EMD-12647), and C9-clusterin focused 3C9-sMAC (EMD-12646). (b) cryoEM density maps (transparent surface) overlaid with atomic models for complement proteins (colored according to protein composition). Representative densities for modeled glycans are shown in the bottom panel.

AGC

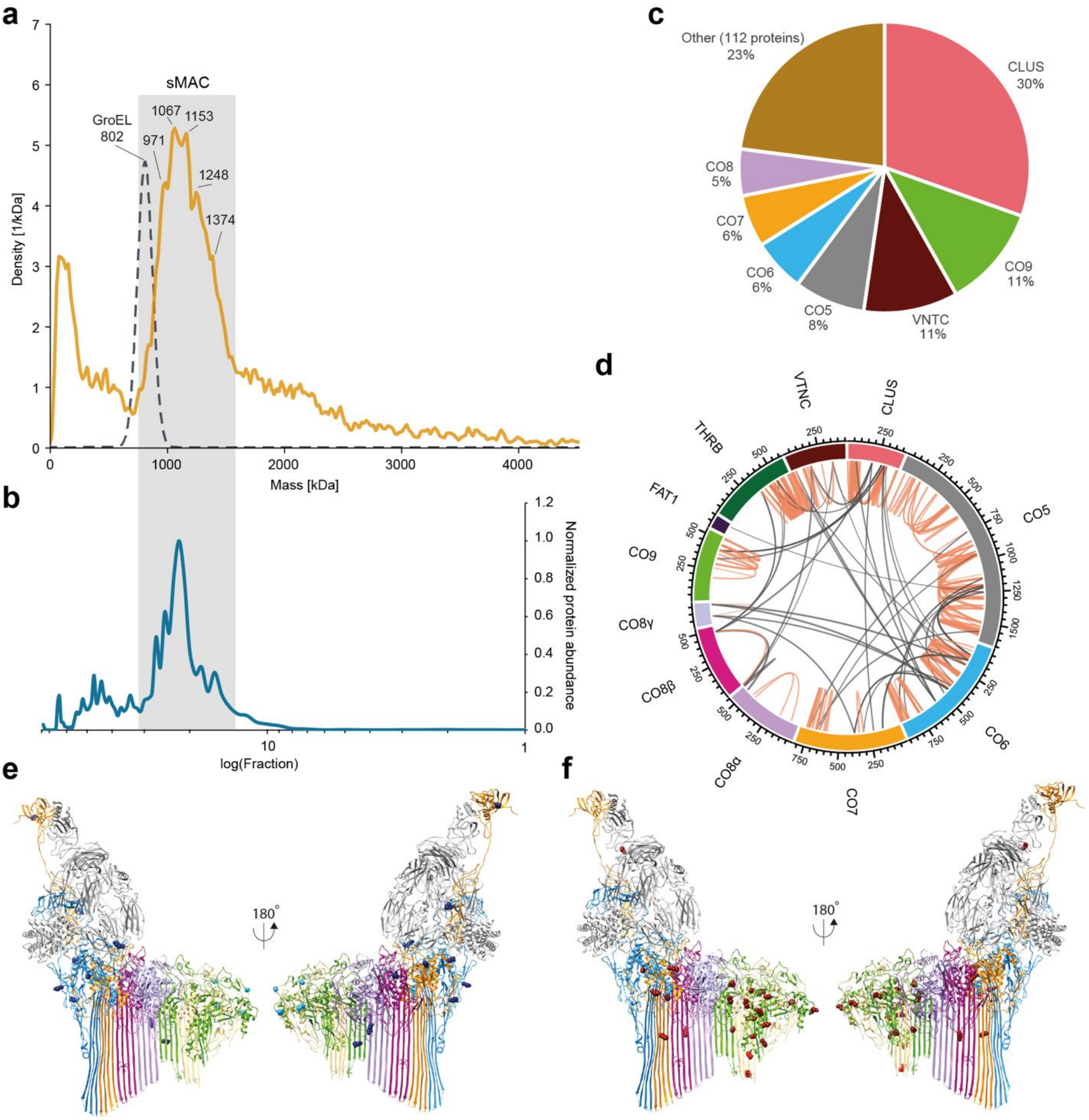


**Supplementary Figure 3:** Single-particle mass photometry and chaperone interactions of sMAC. (a) Mass photometry of sMAC reveals a heterogeneous mixture of complexes. Masses of the most abundant complexes are indicated. Gaussian distribution of GroEL from the protein standard is shown as a reference for peak width of a homogeneous complex. (b) SEC profile of sMAC components (summed abundance) from activated serum. The grey box indicates location of the sMAC complexes in the mass photometry and SEC profile. (c) Label-free quantification (iBAQ) of proteins in the sMAC sample by LC-MS/MS. (d) sMAC circos plot of identified DSS cross-links. The complement components are cross-linked to thrombin (THRB), cadherin 8 domain of protocadherin FAT1 (FAT1) and the chaperones vitronectin (VTNC) and clusterin (CLUS). Intra-links are shown as orange lines and inter-links are shown as black lines. (e) Residues of sMAC uniquely cross-linked to vitronectin are shown as dark blue spheres plotted on the 3C9-sMAC model. Light blue spheres are not resolved in all three C9 molecules. (f) Residues of sMAC uniquely cross-linked to clusterin are shown as dark red spheres plotted on the 3C9-sMAC model. Residues that are not resolved in all three C9 molecules are colored light red. For panels e and f complement proteins are colored as in Fig. 1: C5b (grey); C6 (blue); C7 (orange); C8α (light purple); C8β (dark purple); C8γ (lilac); C9 (alternating green and tan).

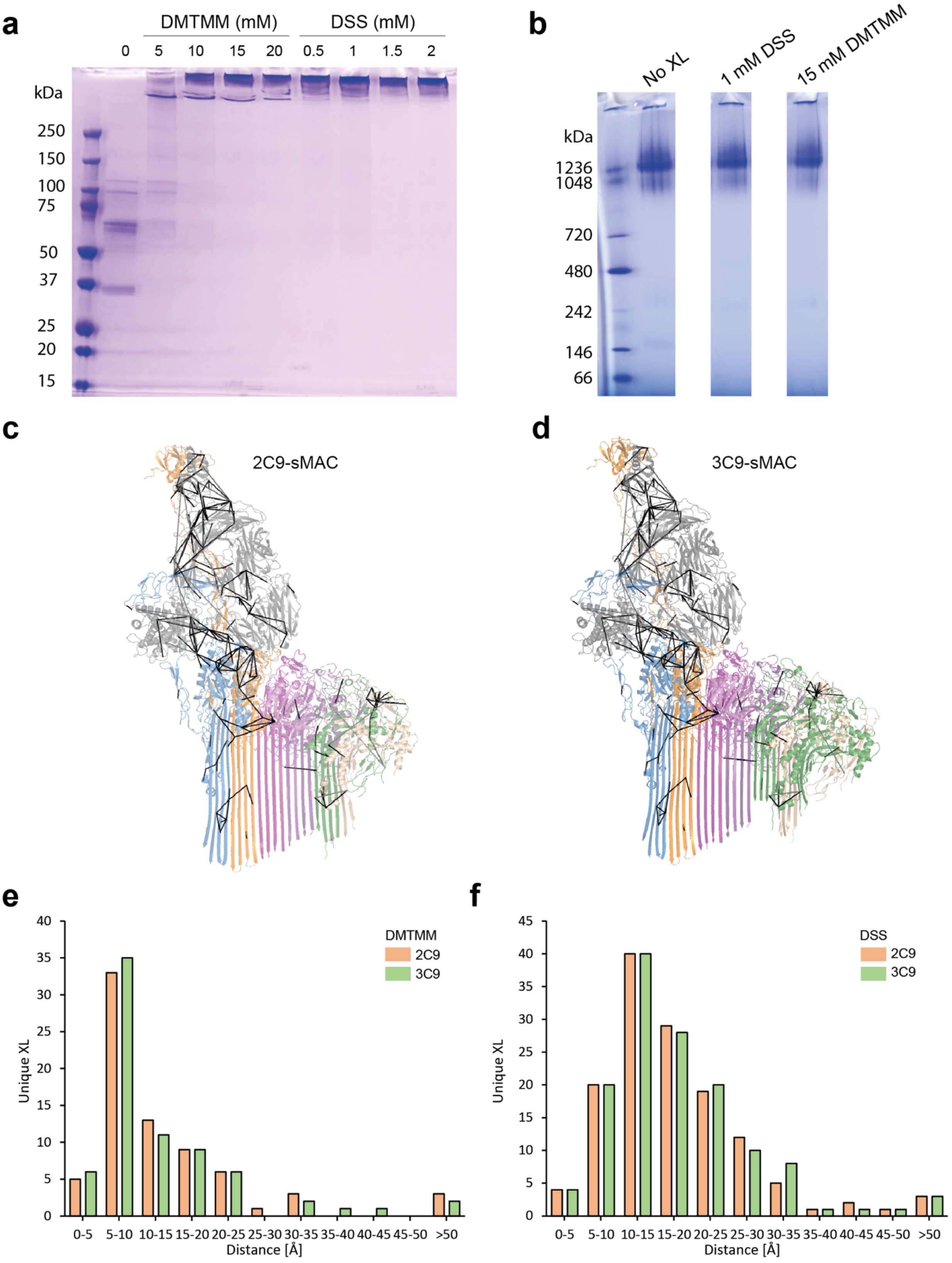


**Supplementary Figure 4:** Cross-linker optimization and mapping of identified cross-links on structural models of sMAC. (a) SDS-PAGE was used to determine optimal cross-linker concentration. sMAC (10 µg) was cross-linked with DMTMM (5-20 mM)

or DSS (0.5-2 mM) and quenched before running on the SDS-PAGE (b) BN-PAGE of cross-linked sMAC (10 µg) using optimal DSS (1 mM) or DMTMM (15 mM) concentrations. (c-d) Identified cross-links plotted on the sMAC structural models containing (c) two C9 or (d) three C9 molecules. Cross-links below the distance restraints (DSS <30Å and DMTMM < 20Å) are shown as black lines and distances above the restraints are shown as dashed grey lines. Complement proteins are colored as in Fig. 1: C5b (grey); C6 (blue); C7 (orange); C8α (light purple); C8β (dark purple); C8γ (lilac); C9 (alternating green and tan). (e-f) Histogram of Cα-Cα distances of (e) DMTMM and (f) DSS cross-links mapped on the 2C9- and 3C9-sMAC atomic models. Displayed cross-links were identified in all three replicates.

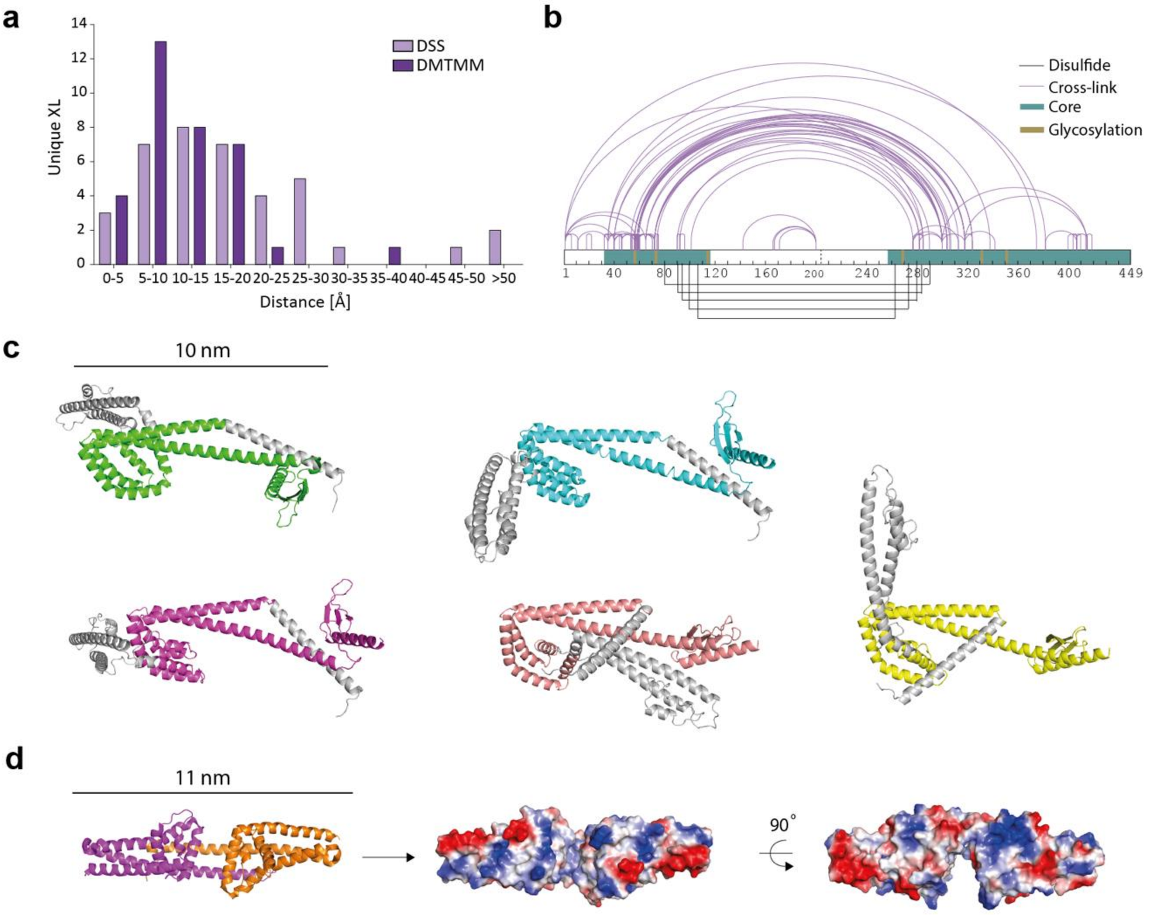


**Supplementary Figure 5:** Structural model for clusterin. (a) Histogram of Cα-Cα distances of DSS and DMTMM cross-links plotted on the clusterin core model. (b)

Identified clusterin inter-links are mapped onto the clusterin sequence shown with purple lines. The five disulfide bridges are shown as black lines. The common core is colored teal and reported glycosylation sites are marked in sand. (c) Panel of structural models generated by trRosetta. The consensus core region is uniquely colored by model, while long helical extensions that flexibly hinge from the core are grey. (e) Structure of CspA from *Borrelia burgdoferi* (PDB ID: 1W33) shown as ribbons (left panel) and electrostatic surface representation (middle and right panels).

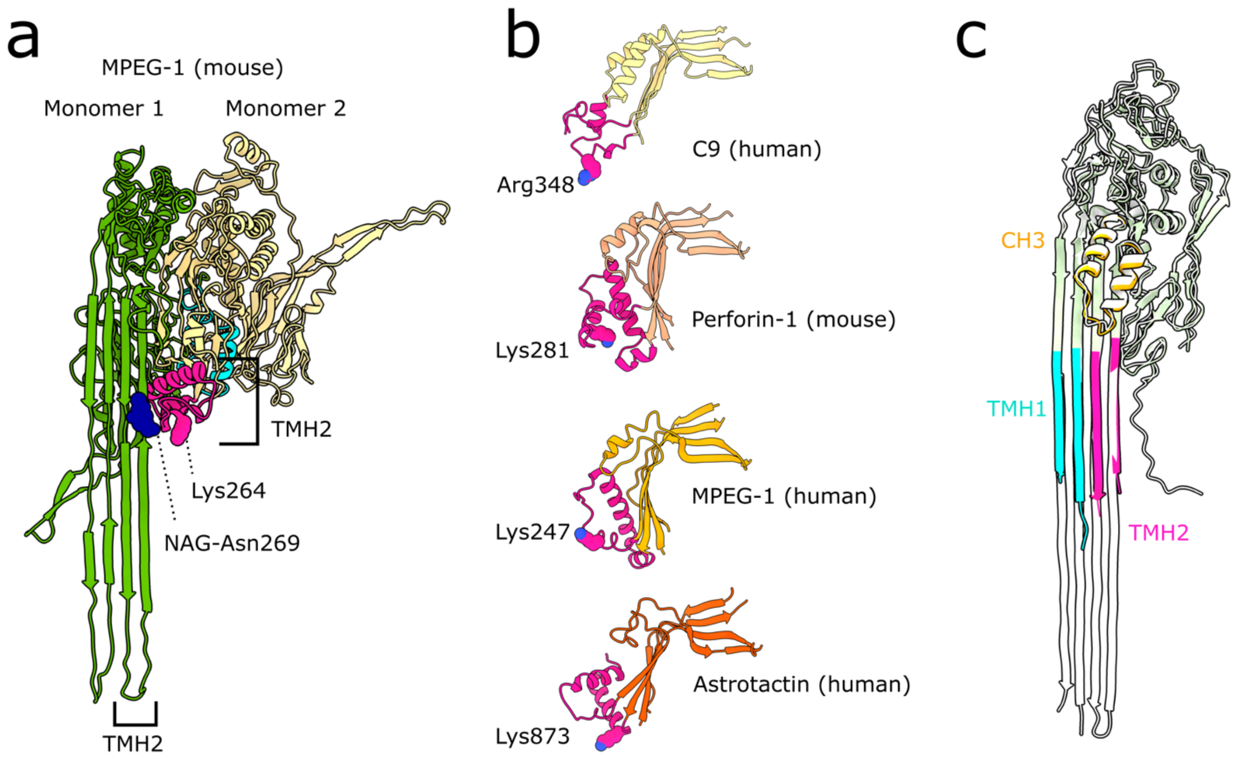


**Supplementary Figure 6:** MACPF structural transitions. (a) Model of a potential intermediate confirmation of MPEG-1 generated by superposing a single monomer from a soluble conformation (PDB ID: 6SB3) onto the MPEG pore structure (PDB ID: 6SB5). Two adjacent MPEG monomers are shown. Monomer 1 (green) is from the transmembrane conformation. Monomer 2 (yellow) is the superposed soluble conformation. This superposition places a positively charged residue (Lys264) within

TMH2 of monomer 2 (pink) near a glyan (blue) on the extended β-hairpins of the preceding monomer (monomer 1, green). TMH1 of the soluble monomer (cyan) is shown for reference. (c) Central kinked β-sheet and TMH2 helical bundle of the MACPF domains of C9 (intermediate conformation in sMAC), peforin-1 (PDB ID: 3NSJ), MPEG-1 (PDB ID: 4OEJ) and astrotactin-2 (PDB ID: 5J68). In each case, a similarly oriented positively charged residue with TMH2 is highlighted (spheres). TMH2s are indicated in pink; TMH1s are removed for clarity. (c) Superposition of the transmembrane conformation of C9 as seen in MAC (white, PDB ID: 6H03) with the penultimate C9 in 2C9-sMAC (colored ribbons: CH3 is yellow; TMH1 is cyan; TMH2 is pink; the remainder of C9 is green).
